# NEUROGLIAL CB1 RECEPTORS CONTROL NAVIGATION STRATEGIES

**DOI:** 10.1101/2025.05.27.656352

**Authors:** Jon Egaña-Huguet, Lucía Sangroniz-Beltrán, Andrés M. Baraibar, Pablo Reyes-Velásquez, Nicolas Landgraf, Paula Torres-Maldonado, Cathaysa Rodriguez-Cedrés, Francisca Julio-Kalajzic, Joaquín Piriz, Almudena Ramos, Pedro Grandes, Giovanni Marsicano, Susana Mato, María Ceprián, Edgar Soria-Gómez

## Abstract

Navigation and memory functions are essential for survival and are regulated by the hippocampus. These processes are tightly controlled, and one of the key modulators involved is the endocannabinoid system, particularly through the cannabinoid receptor type-1 (CB1). CB1 is widely expressed in various hippocampal cell types. While it is known that CB1 participates in memory processes, its specific roles in different cell types and how these roles may differ between sexes remain unclear.

This study investigates how CB1 signaling in the hippocampus, in both, cell-type-specific and sex-dependent manner, contributes to navigation and memory. To this end, we selectively deleted CB1 receptors from neurons, CAMKII-expressing neurons, and astrocytes from the hippocampus of adult male and female mice. We then assessed its effect on a broad range of behaviors, including innate emotional responses, memory, navigation, and other hippocampus-related functions such as nesting.

Deletion of CB1 in CAMKII-expressing neurons produced a pronounced effect in males, leading to increased anxiety and impairments in both reference and spatial memory. These mice also showed altered performance in the Barnes maze, relying less on spatial strategies. By contrast, females were less affected by this specific deletion. Interestingly, only deletion of CB1 from astrocytes led to spatial memory impairments in females, which also showed reduced LTP and a decreased reliance on spatial strategies in the Barnes maze. In conclusion, our findings show that neuronal CB1 receptors are critical for the spatial navigation strategy in males, while astrocytic CB1 receptors play a key role in memory processes both in males and females.

## Introduction

Navigating a landscape is crucial for the survival of species. Animals utilize environmental cues (e.g., visual signals, odors) and previously learned information to achieve specific goals, such as finding shelter or avoiding danger. The strategies employed to perform these behaviors stem from a balance between following spatial cues, flexibility in their interpretation, and efficiently executing experience-guided tasks. In this manner, internal maps are created and can later be utilized to enhance behavioral efficacy (*1–3*). In a novel environment, exploratory behavior may seem disorganized. However, as the environment becomes familiar, vector-based navigation (allocentric strategy) and egocentric strategies typically emerge (*4–7*). Nonetheless, the process of selecting different strategies for specific environments remains unclear.

The hippocampus (HC) (*4–6*), along with other brain regions (*7, 8*), modulates specific strategies and memory formation. Although the hippocampal formation operates as a unified entity in memory processes, the areas along the dorsoventral axis serve distinct roles. The projections received by the HC from cortical or subcortical regions also differ between the dorsal and ventral areas (*9*). The dorsal part primarily processes previously acquired spatial information, while the ventral part plays a more significant role in forming new environmental layouts (*10*).

Different cell types are identified as key components of the navigation process. From place cells in the CA1 region of the hippocampus, which form neural ensembles activated when the animal visits a specific location, to grid cells in the entorhinal cortex, which integrate information about position, direction, and distance (*11*). Furthermore, glial cells, particularly astrocytes, also play a fundamental role in acquiring and utilizing environmental information (*12*, *13*). Accordingly, several neuromodulatory systems regulate these functions, with the endocannabinoid system (ECS) being one of the most studied (*14–16*). In particular, cannabinoid receptor type-1 (CB1) is widely expressed in hippocampal cells, including neurons and glia (*17*, *18*). It is also found in subcellular domains, such as mitochondrial membranes (*19–21*). At the functional level, neuroglial CB1 receptors in the hippocampus are essential for synaptic plasticity and memory formation (*22–24*).

Most of these studies were conducted on male rodents. However, recent research indicates that the role of astrocytes in memory recall and navigation varies between male and female mice (*25*). Stress has also been found to affect spatial learning differently in both sexes, impairing learning only in males (*26–28*). This sexual dimorphism is also evident in endocannabinoid levels (*29*, *30*) and CB1 distribution (*31*, *32*). In this context, little is known about the role of hippocampal CB1s in females, where their involvement in hippocampal-dependent processes appears to differ from that one in males.

In this study, we aim to investigate the cell-specific and sex-dependent role of CB1 signaling in hippocampus-dependent behavioral processes. To achieve this, we have specifically deleted neuronal or astrocytic CB1 in the hippocampus of adult male and female mice and assessed its impact on innate emotional responses, hippocampus-dependent memory and navigation, and synaptic plasticity.

Our tests show that CB1 exclusively impacts the innate emotional response in males. It is also essential for object recognition memory and the development of spatial navigation strategies in male mice. However, loss of CB1 from astrocytes hinders learning and memory recall in both males and females. Finally, CB1 deletion decreased synaptic plasticity in the HC only in male mice.

## Methods

### Animal models

All experimental procedures were approved by the Animal Research Ethical Committee of the University of País Vasco (M20/2019/199 // M30/2019/301). CB_1_-flox animals were a generous gift from Prof. Manuel Guzmán (Complutense University of Madrid). *CB_1_*-KO mice were locally produced at the University of País Vasco Animal Facility. Animals were maintained in individual cages under standard conditions with water and food *ad libitum*.

### Viral vectors

Adenoviral vectors (ssAAV) were purchased from the University of Zurich. The vectors ssAAV-9/2-hSyn1-mCherry_iCre-WPRE-hGHp(A), ssAAV-9/2-hGFAP-mCherry_iCre-WPRE-hGHp(A) and ssAAV-9/2-mCaMKIla-mCherry 2A iCre-WPRE-SV40p(A) were used to selectively deliver Cre expression to neurons, astrocytes and principal glutamatergic neurons, respectively. ssAAV-9/2-hSyn1-chl-mCherry-WPRE-SV40p(A), ssAAV-9/2-hGFAP-mCherry-WPRE-hGHp(A) and ssAAV-1/2-mCaMKIIα-mCherry-WPRE-hGHp(A) were used as control (Table 1).

**Table 1.** ssAAV Viral vectors used in the different experiments.

| ssAAV Viral Vector | Group Name |
| --- | --- |
| ssAAV-9/2-hSyn1-mCherry_iCre-WPRE-hGHp(A) | HC-(Syn)*CB1*-KO |
| ssAAV-9/2-hGFAP-mCherry_iCre-WPRE-hGHp(A) | HC-(GFAP)*CB1*-KO |
| ssAAV-9/2-mCaMKIIα-mCherry 2A iCre-WPRE-SV40p(A) | HC-(CAMKII)*CB1*-KO |
| ssAAV-9/2-hSyn1-chl-mCherry-WPRE-SV40p(A) | CTL |
| ssAAV-9/2-hGFAP-mCherry-WPRE-hGHp(A) | CTL |
| ssAAV-1/2-mCaMKIIα-mCherry-WPRE-hGHp(A) | CTL |

### Surgery and viral vector administration

To delete CB1 in the HC, two-month-old male and female *CB_1_*-flox mice were bilaterally injected in the dorsal hippocampus with the AAVs using the RWD stereotaxic system (RWD; Guangdong, China). The animals were anesthetized with isoflurane at 5% for induction and then maintained at 1% during the surgery. Bilateral injections of 250 nl were carried out at the coordinates −2 AP, ±1.5 ML, and −2 DV, using the Nanoject III system (Drummond Scientific; Broomall, PA, USA). Breathing and body temperature were monitored throughout the procedure. Lidocaine was topically administered at the beginning, and carprofen (4mg/kg) was subcutaneously injected at the end of the surgery. The animals were individually caged with humidified food and observed for a period of three days. Four to five weeks after AAVs hippocampal injection, animals were sacrificed, and brains were processed to verify the site of infection using a fluorescent co-immunostaining for the virus fluorescent protein (mCherry) together with neuronal (NeuN) or astrocytic (GFAP) markers (Fig. S1).

### Behavioral tests

Animals were allowed to recover for four weeks after surgeries to ensure the expression of the adenoviral vector and its effect on disrupting the *de novo* synthesis of the receptor. Behavioral tests were conducted during the light phase, one per day. Animals were allowed to acclimate for 30 minutes in the room before the start of the experiments. Tests were recorded and analyzed using the ANY-maze system (ANY-maze; Dublin, Ireland), except for the novel object recognition test. The animals underwent several behavioral tests in the following order.

### Innate emotional responses

*Open Field*: Animals were positioned in the center of a dimly lit open-field arena (40 × 40 × 45 cm, 200 lux) and allowed to explore freely for 5 minutes. A square area of 6×6 cm, located 6 cm from the walls, was designated as the “center zone,” while the remainder was defined as the “safe zone”. Time spent in each region, traveled distance, and mean speed were analyzed to evaluate anxiety-like behavior. *Elevated Plus Maz*e: Mice were placed in the center of an Elevated Plus Maze (with 4 arms measuring 30×5 cm; 2 open and 2 closed at a height of 49 cm) and allowed to explore the maze freely for 5 minutes. Time spent, number of entries, and traveled distance in both closed and open arms were quantified. *Light-Dark Box*: Mice were placed in a brightly illuminated arena (1200 lux) connected to a dark chamber (each chamber measuring 30×30×35 cm). Time spent, distance traveled, and number of entries into the illuminated area were analyzed.

### Memory

#### Novel Object Recognition Test

The Novel Object Recognition Test (NORT) was conducted using an L maze (with external and internal L walls measuring 35 cm and 30 cm, respectively, and wall dimensions of 4.5cm wide and 15 cm high) (*33*, *34*). All stages were recorded using an iDS camera paired with a uEYE software system. On the first day (habituation), mice were allowed to explore the maze for 9 minutes. On the second day (acquisition), two identical objects were placed at the end of each arm, and mice were given 9 minutes to explore them before returning to their home cage. The test took place 24 hours later, with one of the objects replaced by a new, unfamiliar one. The animal was then allowed to explore both objects for an additional 9 minutes.

Two different treatment-blinded experimenters quantified the time spent exploring each object. The discrimination index (DI) was calculated as the difference between the time spent exploring the novel object (TN) and the familiar object (TF), divided by the total exploration time (TN+TF): DI = [TN-TF]/[TN+TF](*24*). All objects were previously assessed so that animals would not show any preference.

### Memory and navigation

#### Barnes maze

To assess memory and navigation, animals were subjected to the Barnes Maze (BM). The BM is a circular platform of 150cm in diameter, with 20 holes in the vicinity, 90cm above the floor. Animals performed BM for 5 days, four trials per day. To complete each trial, they can use a maximum of 2 minutes, with one resting minute in the home cage between tests. Meanwhile, the maze was cleaned. On the first day, animals are first driven to the target hole before the beginning of the trials. The test begins in the center of the rounded maze, where mice are placed inside a plastic opaque cylinder for 10 seconds and then released. Visual cues are located around the room to ensure the use of the spatial cues to resolve the maze.

The time to reach the target hole, the number of errors, and the strategy used were analyzed using ANY-maze software by a treatment-blind experimenter. Any hole inspected before reaching the target hole is considered an error. Solving strategies were classified as previously described (*5*, *35*). Briefly, when the animal reaches the hole without following any pattern or missing two consecutive holes, it is considered RANDOM. A SERIAL strategy is defined when the animal reaches the hole after exploring consecutively more than one hole with only one missing hole between consecutive holes. Finally, strategy is considered SPATIAL when the animal goes directly to the target or the two nearby holes. If a mouse does not get to the target hole within 5 minutes, it will be gently guided to it. After each trial, mice were left inside the escape box for 1 minute before moving to the home cage.

### Ex-vivo electrophysiology

We collected acute coronal hippocampal slices (350 μm thick) from *CB_1_*-flox mice (males and females) carrying a specific CB1 deletion in hippocampal neurons (Syn) or astrocytes (GFAP). Slices were kept in ice-cold artificial CSF (ACSF) containing the following (in mM): 124 NaCl, 5 KCl, 1.25 NaH_2_PO_4_, 2 MgSO_4_, 26 NaHCO_3_, 2 CaCl_2_, and 10 glucose gassed with 95% O_2_/5% CO_2_, pH 7.3–7.4. Slices were incubated in ACSF at room temperature for at least 1 h before use, then transferred to an immersion recording chamber, superfused at 2 mL/min with gassed ACSF, and visualized under an Olympus BX51WI microscope (Olympus Optical, Japan). Field excitatory postsynaptic potentials (fEPSPs) were evoked in the CA1 stratum radiatum by stimulating Schaffer collaterals (SCs) with theta capillaries (2–5 µm tip) filled with ACSF and recorded with ACSF-filled glass pipettes (< 5 MΩ). Electrical pulses were supplied by a stimulus isolation unit (ISO-Flex, A.M.P.I., Jerusalem, Israel). Signals were fed to a Pentium-based PC through a DigiData 1550 interface board. Signals were filtered at 1 kHz and acquired at a 10 kHz sampling rate using a DigiData 1550 data acquisition system. pCLAMP 10.3 software (Molecular Devices, San Jose, CA) was used for stimulus generation, data display, acquisition, and storage. For baseline recording, the stimulation intensity was adjusted to obtain 40% of the maximum slope of the response, and inputs were stimulated (1 ms pulse duration) every 5 s. The slope of the fEPSPs was measured between 30 and 70% of the maximum. For LTP induction, a tetanic stimulation (4 trains at 100 Hz for 1 s; 30 s intervals) was applied to the SCs. fEPSP slope was normalized to the 10 min of baseline recording. After LTP induction, fEPSPs were recorded for 60 min. The presence of LTP was determined by comparing the average fEPSP slope during the baseline with the mean values from the 10–30 min (early LTP) and 30–60 min (late LTP) post-induction periods.

### Tissue processing

Once behavioral tests were performed, mice were deeply anesthetized by an intraperitoneal administration of a mixture of Ketamine/Xylazine (80/10 mg/kg body weight). Right after, they were transcardially perfused with PBS (0.1M, pH 7.4) for 1 min. before being fixed with 4% formaldehyde solution for another 7 minutes with a peristaltic pump at 12ml/min (Perimax 12; Spetec GmbH, Erding, Germany). Brains were extracted and post-fixed in a 4% formaldehyde for one week at 4°C. For their storage, the fixative solution was diluted 1:10 and brains were kept at 4°C until use.

### Immunofluorescence

Stored brains were cut into 50µm slices using a vibratome (Leica, VT1000S) and kept in PB 0.1M + azyde 0.05%. To analyze the depletion of the CB1 receptor, slices from AP - 1.7 to AP −2.3 were selected. Slices were washed with PB 0.1M to clean the azyde and then blocked for 1 hour at room temperature (RT) with blocking buffer (PB 0.1M + donkey normal serum at 10% (S30M, Sigma-Aldrich; St. Louis, USA) + 0.3% triton X-100 (X100-5ML, Sigma-Aldrich; St. Louis, USA). Slices were then incubated with primary antibodies for 72 hours at 4°C. To verify the viral vector injection site, together with the co-localization in astrocytes, several antibodies were used: mouse anti-Hexaribonucleotide Binding Protein-3 (α-NeuN; RRID_AB104224, Abcam; Cambridge, UK) at 1:1000; rabbit anti-Cherry monomer (α-mCherry, RRID_AB356482, Abcam; Cambridge, UK) at 1:500; mouse anti-glial fibrillary astrocytic protein (α-GFAP; RRID_AB257130, Abcam; Cambridge, UK) 1:1000. Slices were washed with PB 0.1M before adding secondary antibodies for 3 hours at RT. The secondary antibodies were either α-rabbit AlexaFluor 594 (RRID_AB2534016), α-mouse AlexaFluor 488 (RRID_AB141607). Finally, mouse brain slices were washed and mounted using Fluoroshield Mounting Medium with DAPI (F6057; Sigma-Adrich; St. Louis, USA). Microphotographs from hippocampi were taken using a Nikon TiU microscope with a Nikon Ds-Qi2 camera or in a Zeiss Apotome microscope attached to a Axiocam 506 mono y ER5c camera. Images were analyzed using FIJI software (*36*).

### Nest building

Nests were collected every 2 weeks. Nest pictures were obtained using a black cardboard with a ruler and a camera (Luminex Panasonic, DMC-FZ200; Panasonic Spain, Cornellá de Llobregat, Spain). The percentage of remaining tissue versus the intact full paper is calculated using FIJI to analyze the degree of nesting complexity (adaptation of Deacon 2006) (*37*). The images are transformed to 8-bit and the threshold is set to 125. The total area of the remaining tissue is calculated and normalized to the original paper size.

### Statistical analysis

For the statistical analysis, we used GraphPad 9 software (GraphPad Prism version 9.0.0 for Windows, GraphPad Software, Boston, Massachusetts USA, www.graphpad.com). First, data normality was assessed using Shapiro–Wilk and Kolmogorov-Smirnov tests. For innate emotional response and NORT tests, an ordinary One-Way ANOVA test was applied, followed by a post-hoc Holm-Sidak’s multiple comparisons test.

BM results were analyzed and compared as the daily mean for each trial. Additionally, the 5-day mean was used to compare genotypes based on latency, number of errors, and strategies. A two-way ANOVA was performed, followed by Tukey’s multiple comparisons test. Electrophysiology *ex vivo* was characterized by a two-tailed unpaired Student *t* test. When data did not meet normality, a Mann-Whitney test was applied.

Results are expressed as mean ± SEM. Significant results were illustrated on the graphs by an asterisk or hashtag, depending on the comparison, representing the different critical P values * p < 0.05; ** p < 0.01; *** p < 0.001; **** p < 0.0001. The results were considered significant when the p-value was equal to or lower than 0.05.

## Results

### Hippocampal CB1 deletion impacts innate emotional response in males, but not females

Four weeks after the injection, the animals underwent a series of behavioral tests (open field, elevated plus maze, and light-dark box) to evaluate their innate emotional response to various aversive stimuli, ranging from the least to the most stressful.

Receptor deletion had different effects on the genotypes and sexes assessed. Differences were mainly found in male mice, in the number of entries and time spent in the center of the open field (Fig.1A, blue charts; F:5.534;p=0.0010 and F:13.19; p<0.0001), in the entries in the open arms of the elevated plus maze (Fig. 1B, blue charts; F:5.993;p=0.0005) and in the time in the light zone of the L-D box (Fig. 1C, blue charts; F:4.570; p=0.0036).

**Figure 1.**
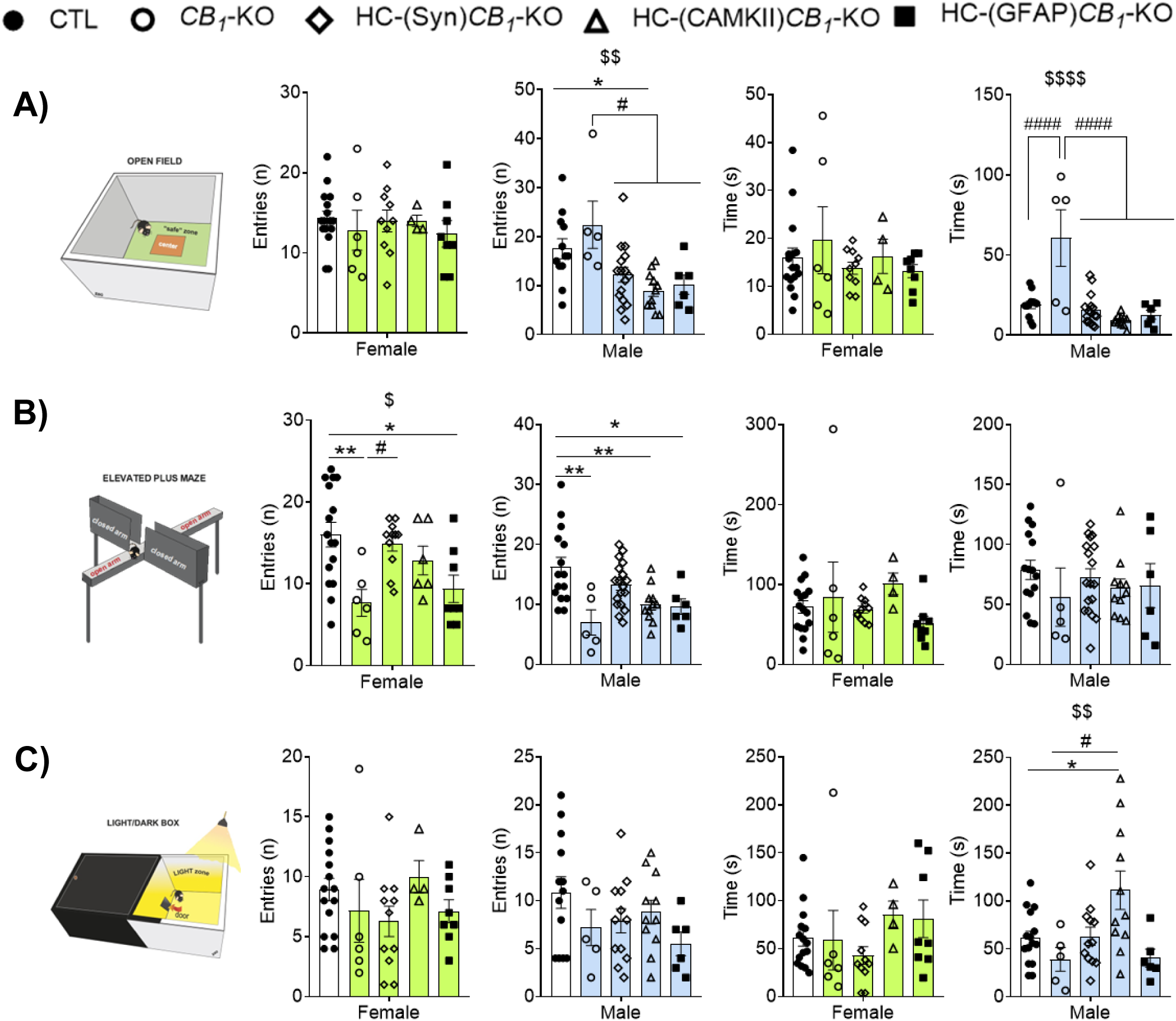
Effect of cellular *CB_1_*-KO in the hippocampus on the innate emotional behavior in male and female mice. Male animals with CB1 deletion in neurons and astrocytes showed an anxiogenic profile with lower entries in the center of the openfield center **(A)** and in the open arms of the elevated plus maze **(B)** and an increased time in the light zone and light and dark box **(C)** compared with control animals. Differences in females were only found in the number of entries to the open arms for the (GFAP)*CB_1_*-KO **(B)**. Data are presented as mean ± S.E.M. $$ p<0.01, $$$$ p<0.0001 differences in One-Way ANOVA test; * p<0.05, ** p<0.01, **** p<0.0001 vs CTL; and # p<0.05, #### p<0.0001 vs *CB_1_*-KO by Holm-Sidak’s multiple comparison tests. See **Methods** for detailed statistics.

In males, full knockout mice (*CB_1_*-KO) showed an increased time spent in the center zone of the open field test (Fig. 1; **** p= <0.0001) and a significant decrease in the number of entries to open arms in elevated plus maze test (Fig. 1; ** p = 0.0024). Besides *CB_1_*-KO, the most affected phenotype was observed when CB1 was specifically deleted from principal neurons ((CAMKII)*CB_1_*-KO) in the hippocampus. HC-(CAMKII)*CB_1_*-KO male mice exhibited a significant decrease in the number of entries in the center zone of the open field test (Fig. 1A; * p = 0.022) and the open arms of the elevated plus maze (Fig. 1B; ** p = 0.0093), accompanied by an increase in the time spent in the light zone of the L-D box (Fig. 1C; * p = 0.0266) compared to controls. Differences were also found when we compared HC-(CAMKII)*CB_1_*-KO performance with *CB_1_*-KO animals, that showed lower number of entries and less time spent in the center of the open field (Fig. 1A; ** p = 0.0033; **** p < 0.0001) and the light zone of the L-D box (Fig. 1C; * p = 0.0178).

Regarding the other genotypes assessed, a significant decrease in the number of entries into the open arm was noted for HC-(GFAP)*CB_1_*-KO male mice when compared to control animals (Fig. 1; * p = 0.0317) and in the entries and time in the entries and time when compared to *CB_1_*-KO animals (Fig. 1A; * p = 0.0234; **** p < 0.0001).

In female mice, CB1 deletion did not induce a clear emotional alteration, neither in the global KO nor in the cell-specific KO, with the only differences observed in the elevated plus maze test for the *CB_1_*-KO (Fig. 1B; ** p = 0.0077) and HC-(GFAP)*CB_1_*-KO (Fig. 1; * p = 0.024) female genotypes.

Taken together, our results show that the deletion of CB1 alters the innate emotional response in males but not in females, with the deletion of CB1 from principal neurons having a significant impact on the assessed behavior.

### Hippocampal CB1 deletion from neurons and astrocytes impaired memory in males and females, respectively

To address the effect of sex and cellular deletion of CB1 on memory processes, we used the novel object recognition and Barnes maze (BM) tests. To assess recognition memory, we performed the NOR test in the L maze. CB1 deletion, either global or cell-specific, has no effect on the D.I. in females but has a negative impact (Fig. 2; F = 1.820; p = 0.1717) on the memory evaluated in males (Fig. 2; F = 8.509; #### p < 0.0001). A significant reduction in the D.I. was observed in males in which CB1 was deleted from CAMKII-positive cells (Fig. 2B; ** p=0.0056), which also showed a significant decrease in the D.I. compared to full knockouts (Fig. 2B; * p = 0.0305). A similar trend was observed in HC-(CAMKII)*CB_1_*-KO (Fig. 2B; * p = 0.0716). In the BM, mice learn the position of the hidden escape box, using different visual cues located in the behavioral room, with the test repeated for 5 days. A significant memory impairment in both sexes was observed, with a substantial impact on the latency to enter the scape box (Fig. 3; Females: F: (4, 45) = 6.785; ### p = 0.0002 // Males: F (3, 51) = 4.201; ## p = 0.0099), and in the number of errors committed before finding the scape box (Fig. 3; Females: F (4, 43) = 5.344; ## p = 0.0014 // Males: F (4, 53) = 3.188; # p = 0.0203). Female HC-(GFAP)*CB_1_*-KO mice showed the biggest increase in the latency to reach the target hole (Fig. 3A; ** p = 0.0021) and an increase in the number of errors (Fig. 3B; * p = 0.0299), compared to control mice. By contrast, HC-(Syn)*CB_1_*-KO and HC-(CAMKII)*CB_1_*-KO male mice presented an increase in latency and errors (Fig. 3; **** p < 0.0001; *** p = 0.0006) (Fig. 3; **** p < 0.0001; ** p = 0.0029) groups affected.

**Figure 2.**
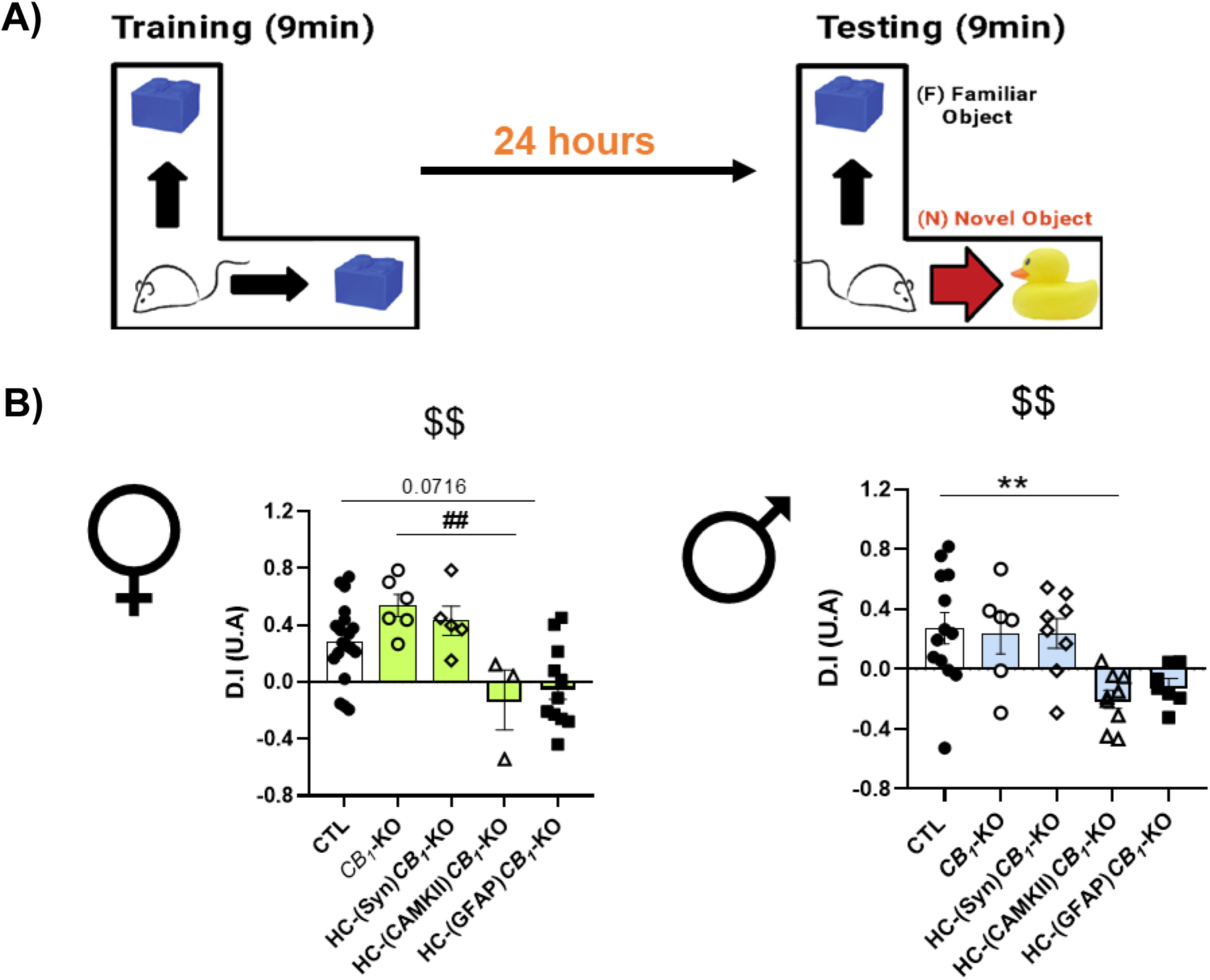
CB1 deletion in CAMK neurons of the hippocampus impairs object memory only in male mice. **(A)** Animals were introduced to two identical objects for 9 minutes for training. 24 hours later, the animals were re-exposed, and one of the objects was changed for a new one. In females, CB1 deletion in glutamatergic neurons induced a lower D.I when compared to global *CB_1_*-KO and showed a decreased trend when compared to controls (p=0.0716) **(b, left panel)**. By contrast, (CAMK)*CB_1_*-KO severely impaired male object recognition memory compared to control **(b, right panel)**. Data are presented as mean ± S.E.M. #### p<0.0001 differences in One-Way ANOVA test; ** p<0.01 vs CTL; ## p<0.01 vs *CB_1_*-KO by Holm-Sidak’s multiple comparison tests. See **Methods** for detailed statistics.

**Figure 3.**
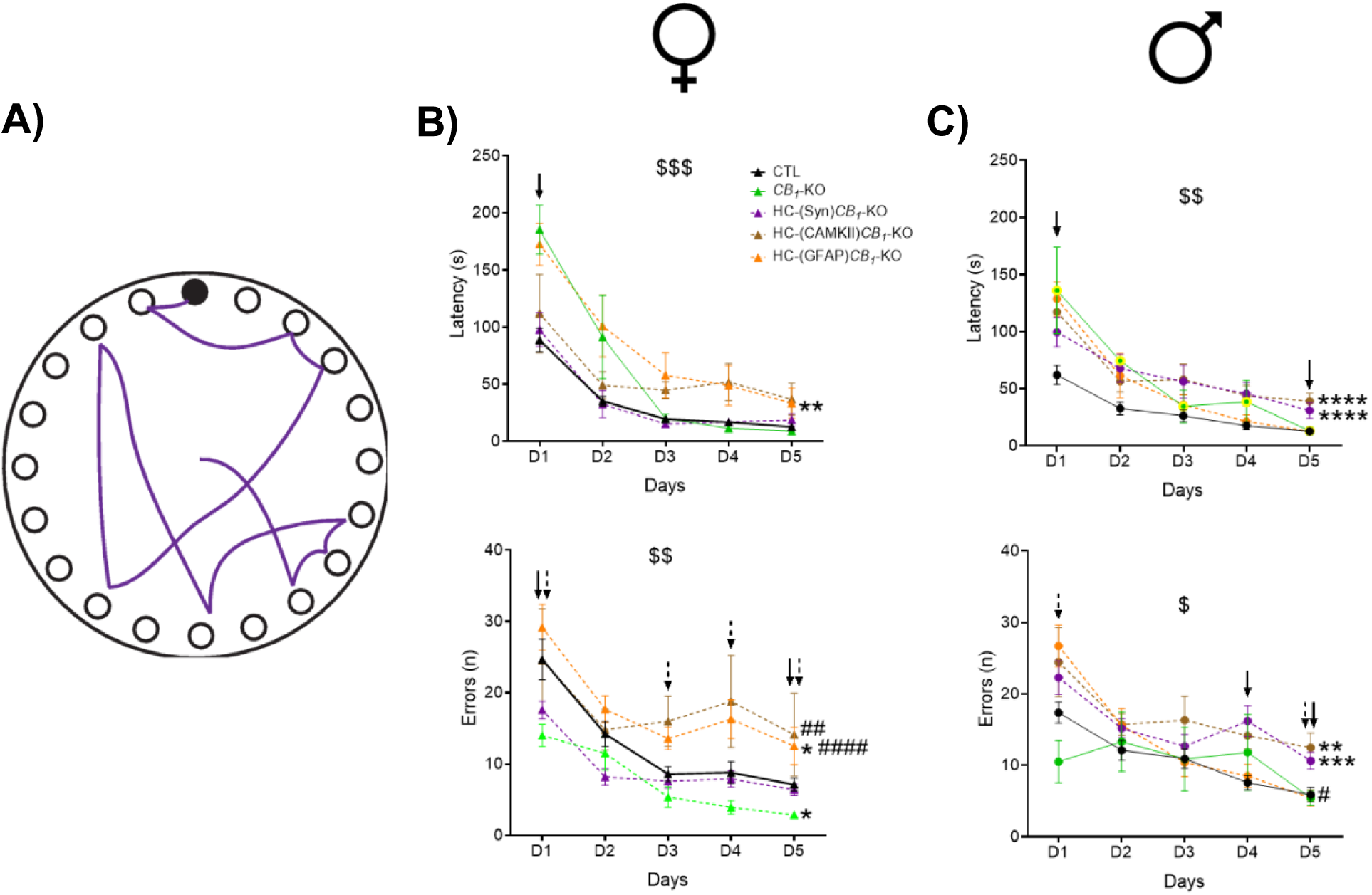
Sexual dimorphism on CB1 deletion impact on navigation strategies. **(A)** Animals performed the Barnes maze for five consecutive days, and the latency and number of errors made before reaching the target hole were recorded. Females with the astrocytic CB1 deletion showed a clear learning impairment, needing more time to reach the target hole **(B, left panel)**. The number of errors was also increased **(C, left panel)**, with significant differences per day with the control (***black arrow***) and the CBN **(*dashed arrow***) groups. By contrast, significant differences were found in males with CB1 deletion in neurons (HC(Syn)*CB_1_*-KO) and glutamatergic neurons (HC(CAMK)*CB_1_*-KO) in both the latency to reach the target hole **(B, right panel)** and in the number of errors made **(C, right panel)**. Day differences with **→** the control (***black arrow***) and the CBN **(*dashed arrow***) animals were also found. Data are presented as mean ± S.E.M. $ p<0.05, $$ p<0.01, $$$ p<0.001 differences in Two-Way ANOVA test. * p<0.05, ** p<0.01, **** p<0.0001 vs CTL; ## p<0.01, #### p<0.0001 vs CBN, p<0.05 vs CTL; p<0.05 vs CBN. Females’ results were F(4, 45)=6.785; $$$ p = 0.0002 for latency and F(4, 43)=5.344; $$ p = 0.0014 for the number of errors. Males’ results were F(3, 51)=4.201; $$ p = 0.0099 for latency and F(4, 53)=3.188; $ p = 0.0203 for the number of errors. See **Methods** for detailed statistics.

Overall, we concluded that CB1 deletion from CAMKII-positive cells in the HC of males negatively impacted both recognition and memory in the BM. By contrast, females’ memory processes were impaired exclusively when CB1 was deleted from astrocytes.

### Specific HC-CB1 deletion impairs the development of spatial strategy in males but not in females

Animals use a wide range of navigation strategies to reach the escape box. In this work, we evaluated three different strategies, spatial, serial, and random; with the spatial strategy being more related to the hippocampal circuitry. In this regard, we found a significant effect on the overall use of spatial strategies in both sexes (Fig. 4; Females: F (4, 38) = 4.418; ## p = 0.005// Males: F (4, 47) = 11.21; #### p < 0.0001) but with a clear sexual dimorphism. In the case of male mice, there was a decrease in the use of spatial strategy after CB1 was deleted from all neurons (Fig.4B; **** p < 0.0001), astrocytes (Fig.4B; * p = 0.043) and glutamatergic principal neurons (Fig.4B; **** p < 0.0001). In female mice, however, the use of spatial strategy was only significantly reduced in those mice where CB1 was deleted from astrocytes (Fig. 4A; ** p =0.0032).

**Figure 4.**
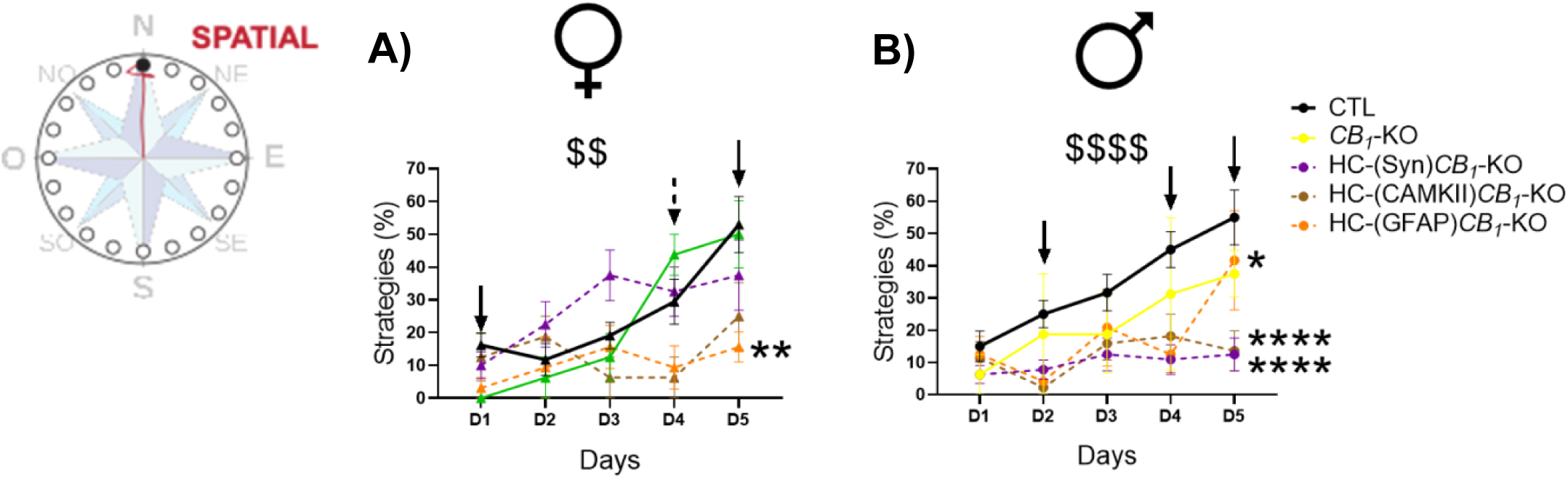
CB1 deletion in the hippocampus impairs spatial strategy acquisition. Over the test days, animals develop a spatial strategy in which they go directly to the target hole using the visual cues **(A)**. The percentage of females with astrocytic CB1 deletion to develop spatial strategies was significantly lower than control mice **(B)**. In males, the acquisition of spatial strategies was impaired by CB1 deletion independently of the cell (neuronal, glutamatergic, or astrocytes) except in the full *CB_1_*-KO **(C)**. Data are presented as mean ± S.E.M. $$ p<0.01, $$$$ p<0.0001 differences in Two-Way ANOVA test. * p<0.05, ** p<0.01, **** p<0.0001 vs CTL, p<0.05 vs CTL; p<0.05 vs CBN. Females’ results were F(4,38)=4.418; ## p = 0.005 and males’ results were F(4, 47)=11.21; #### p = 0.0001. See **Methods** for detailed statistics.

Regarding the other types of strategies, the use of the serial was only reduced in female *CB_1_*-KO mice (Fig. S2; * p = 0.0172), despite no difference being found when the performance of all groups was compared. By contrast, the random strategy use was changed in males but not in female mice (Fig. S2; Random strategy: Females: F (4, 38) = 1,987; ns p = 0.1161// Males: F (4, 48) = 3,589; # p = 0.0122), with a higher percentage of random strategy after deleting CB1 from all neurons (Fig. S2; HC-(Syn)*CB_1_*-KO ** p = 0.0055), from principal neurons (Fig. S2; HC-(CAMKII)*CB_1_*-KO * p = 0.0032) and from astrocytes (Fig. S2; HC-(GFAP)*CB_1_*-KO ** p = 0.0405).

In summary, neuronal CB1 deletion in the HC of male mice negatively impacted the use of spatial strategy with a concomitant increase of the random one.

### Nesting building is differently impaired in males and females

We analyzed the complexity of animals’ nests one month after the surgery to study whether CB1 deletion could impact other complex tasks dependent on HC. Interestingly, the results showed a dimorphism pattern opposite to the one observed in the emotional and memory tests. Males with CB1 astrocyte deletion showed that the nesting paper has been worked on more by the mice when compared to controls (Fig. 5B, p<0.0001), with no difference observed in the neuronal groups. By contrast, only the HC-(CAMKII)CB1-KO females showed an increased level of paper buildout (Fig. 5A, p=0.0002).

**Figure 5.**
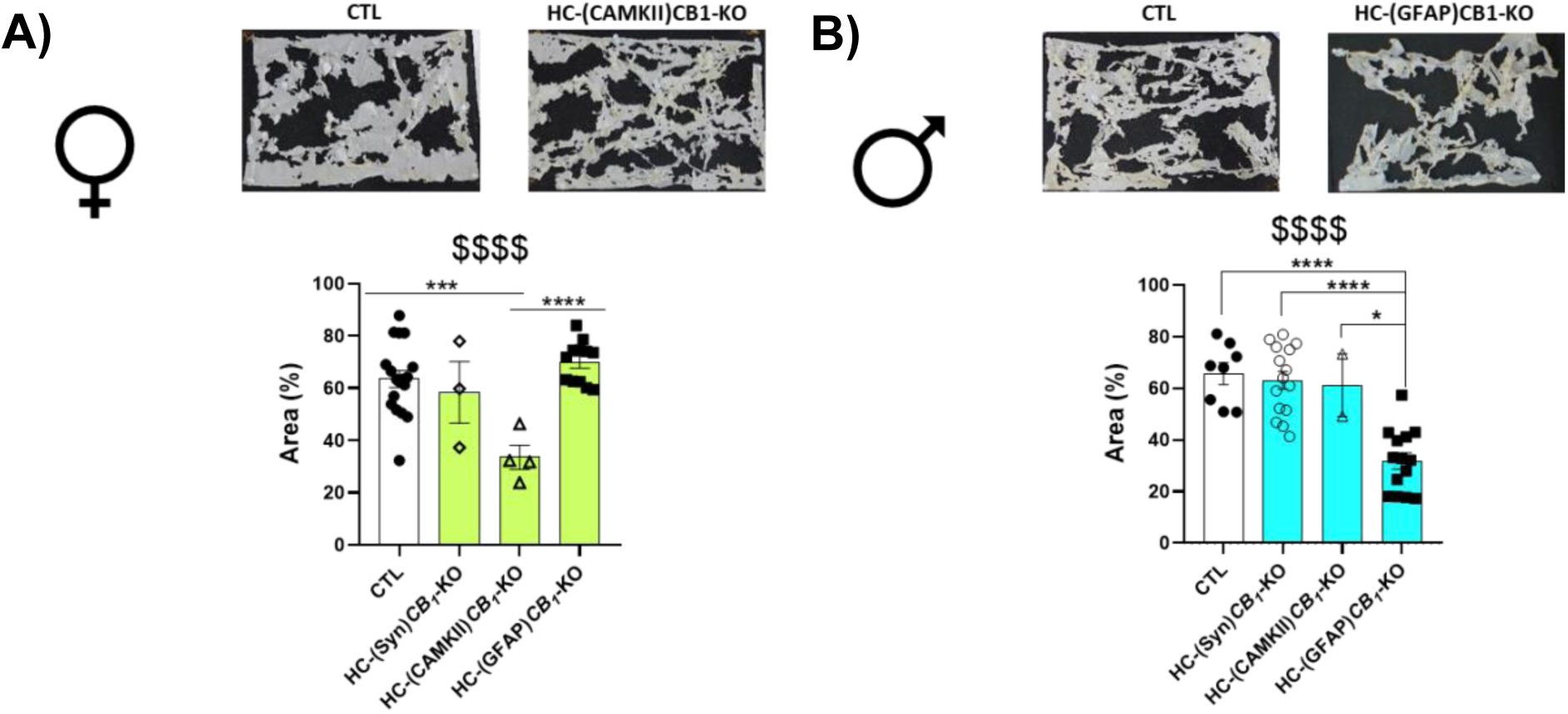
Nesting building is differentially altered in males and females depending on CB1-cellular deletion. Analysis of 2-week-old nests showed that **(A)** females with CB1 deletion in principal neurons (HC(CaMKII)*CB_1_*-KO, triangles) promoted paper nest working compared to control. **(B)** Nest analysis in male animals showed significant differences for HC(GFAP)*CB_1_*-KO group, with a lower percentage of intact remaining nest paper compared to all other neuronal groups and controls. Data are presented as mean ± S.E.M. $$$$ p<0.0001 differences in One-Way ANOVA test; *p<0.05, *** p<0.001, **** p<0.0001 vs CTL; and # p<0.05, #### p<0.0001 vs *CB_1_*-KO by Holm-Sidak’s multiple comparison tests. See Methods for detailed statistics.

### Specific **CB_1_** deletion from neurons and astrocytes in the HC impairs LTP induction in male but not in female mice

To assess the possible mechanism of the impact on memory formation and navigation induced by cellular CB1 deletion, we measured synaptic alteration in the dorsal HC by “*ex vivo*” electrophysiology (Fig. 6A). Field excitatory postsynaptic potential responses (fEPSPs) were measured in CA1 synapses after an HFS protocol was applied in the CA3 subfield of the hippocampus. The results obtained indicate that the LTP induction was significantly impaired in those male and female mice where CB1 was deleted in astrocytes (Fig. 6B; females * p = 0.0411, early LTP and * p = 0.0.0129, late LTP / males * p = 0.0111 early LTP and ** p = 0.0.0068 late LTO). Interestingly, neuronal CB1 deletion only decreased LTP in males, with no differences found in female mice when compared to control animals (Fig. 6C; females n.s. p=0.3196, early LTP and n.s. p=0.6350, late LTP/ males * p=0.0435, early LTP and n.s. p=0.2919, late LTP)

**Figure 6.**
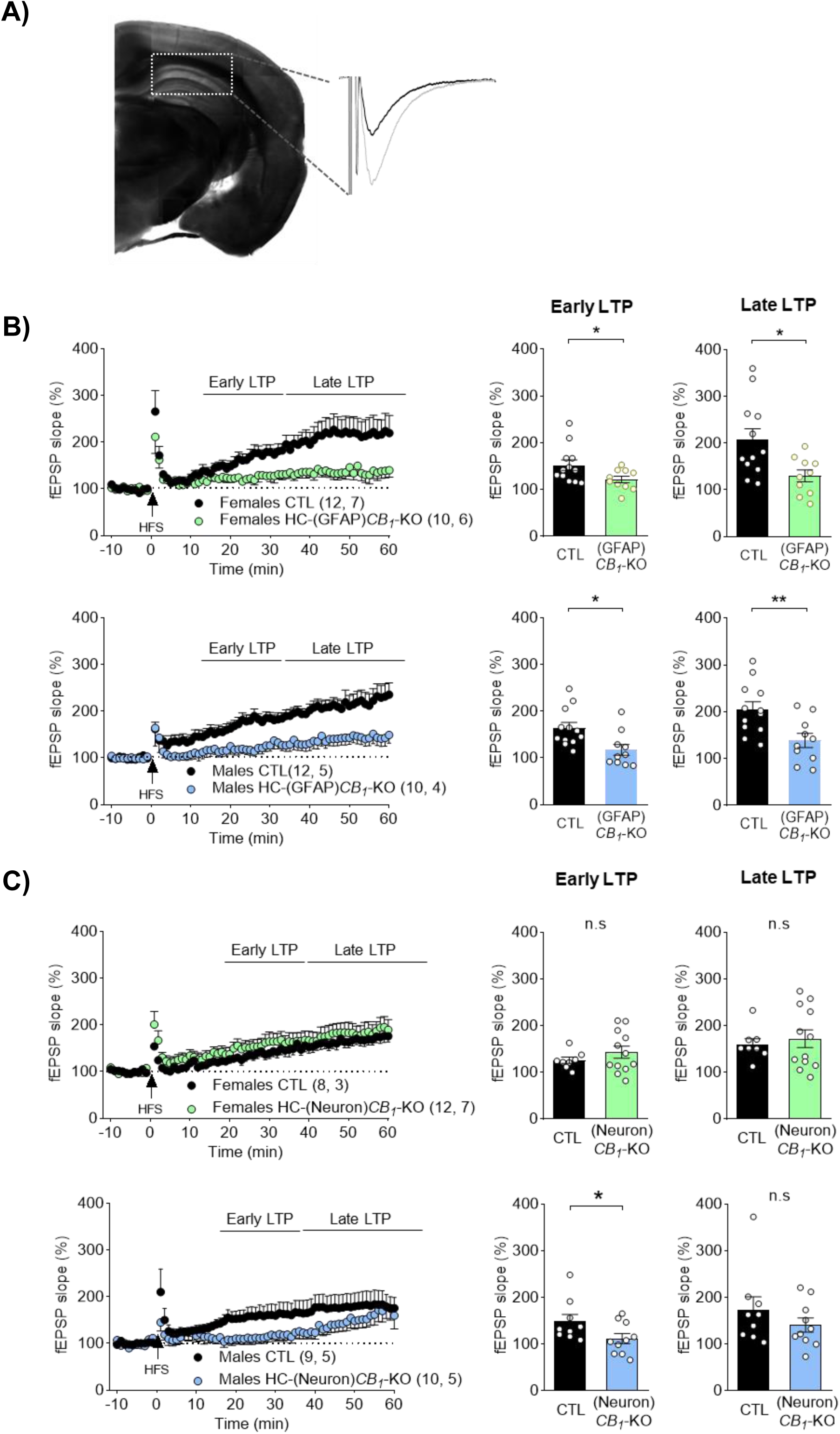
Synaptic plasticity is differentially altered in males and females depending on CB1-cellular deletion. Field excitatory postsynaptic potential responses (fEPSPs) were measured in CA1 synapses after an HFS protocol was applied in the CA3 subfield of the hippocampus **(A)**. CB1 deletion in GFAP astrocytes (HC-(GFAP)*CB_1_*-KO) induced an LTP impairment in both female-upper panel- and male-lower panel-animals. By contrast, CB1 deletion in neurons, including both HC-(Syn)*CB_1_*-KO and HC-(CAMKII)*CB_1_*-KO, induced only an early LTP reduction in males-lower panel-but not females-upper panel-**(C)**. Data are presented as mean ± S.E.M.; *p<0.05; **p<0.001 by two-tailed unpaired Student *t* test. See Methods for detailed statistics.

## Discussion

Previous studies have shown that CB1 pharmacological manipulation impacts referential and spatial memory. However, the specific cells involved in these functions remain unknown, along with the long-standing question of whether these mechanisms are sex-dependent. The current work aims to elucidate the cell-type and sex-dependent role of hippocampal CB1 receptors in memory and navigation. Through this study, we observed that male mice were consistently more affected by CB1 deletion in neurons, particularly in CaMKII-expressing neurons, while females exhibited a milder phenotype, primarily linked to astroglial deletion.

The exception to this observation was the nesting building results. Nest construction is widespread throughout the animal kingdom. They facilitate reproduction, shelter from predators, hibernation, including heat conservation, especially for small-sized rodents. Both male and female mice build nests of similar size, but female nest construction is influenced by hormonal and reproductive status, and nesting ability influences reproductive success (*37*). Animals with hippocampal lesions show an impaired nesting behavior, and the degree of hippocampal lesions is correlated with the impaired level of nest building (*38*) and is affected in hippocampal-related diseases like Alzheimer’s Disease (*39*). However, the cells and mechanisms involved in this crucial function of the hippocampus remain unknown. Our results showed that hippocampal CB1 is involved in nest building, and its deletion led to nesting abnormalities in both males and females; however, the cells involved in this process were different in each sex: CB1 astrocytic deletion in males and CB1 in CAMKII neurons in females. Even though different cells were involved in this, the outcomes were similar, more work was done on the nesting paper. These results were also not linked to increased anxiety levels, since both phenotypes had no differences from controls in the three tests performed. Interestingly, HC-(GFAP)*CB_1_*-KO males had reduced fESP slope, compared to controls, a synaptic disruption that could be related to nesting building impairment without affecting memory, navigation, or the innate emotional state of the animal. By contrast, HC-(GFAP)*CB_1_*-KO females also showed lower fESP percentage, but this did not correlate with nesting building, but it did with navigation and memory. To our knowledge, this is the first time that this has been described, and further studies are needed to assess how hippocampal astrocytes are involved in this complex function.

While astrocytic CB1 deletion in males did not affect the innate emotional behavior of the animals, male HC-(CAMKII)*CB_1_*-KO mice were characterized by anxiety-like behavior through the three tests performed. These animals had lower entries to the center in the OF and the EPM but higher times in the light zone of the light and dark box. These results are aligned with recent studies that point out the role of CAMKII neurons of the ventral hippocampus in anxiety and fear modulation, with their inhibition leading to attenuated anxiety-like behavior (*40*, *41*). Besides this neuronal role, hippocampal astrocytes also respond to anxiogenic environments, and their activation through optogenetics similarly leads to decreased anxiety levels (*42*). In this sense, we could not observe any clear differences in HC-(GFAP)*CB_1_*-KO, but both male and female mice had lower entries in the open arms of the elevated plus maze.

Astrocytic CB1 located in the HC is necessary for learning processes and recognition memory, in both male and female mice, as we observed in the D.I. of the NOR test, and in the latency curve and number of errors of the BM test. This recognition memory impairment was also described in full GFAP-*CB_1_*-KO mice(*22*) and the participation of astrocytic CB1 within the HC has also been described in working memory(*43*) (Shang et al., 2023). Further studies are needed to discern the underlying mechanism. One hypothesis involved the CB1’s regulation of neuronal NMDAR activity through D-serine availability and, thus, modifying LTP(*22*). In agreement with this, our results proved that the lack of astroglial CB1 in HC was sufficient to induce LTP decrease in both sexes, although it was only related to memory impairment in females. Besides this mechanism, activation of astrocytic CB1 also upregulates glutamatergic transmission through different gliotransmitter releases like the eCBs, which participate in the tripartite synapse(*44*). The fact that GFAP is also expressed in hippocampal progenitor cells should also be considered. Astrocytic CB1 promotes astrocyte differentiation through endocannabinoid signaling(*45*). Its manipulation may alter astrocyte-driven functions (*46*), potentially affecting neurogenesis, learning, and memory formation(*47*).

In the BM, neuronal CB1 receptors are necessary in males since their deletion increases both the latency and the number of errors in this test. Navigation is a complex task that requires the animal to identify and learn the objective position, and then navigate toward it using internal (egocentric) or external (allocentric) cues. BM results showed that spatial navigation was significantly reduced in males after CB1 was deleted from hippocampal neurons, a deletion that did not affect females. This could be partly explained by the fact that female mice do not use this strategy as frequently as males. In our study, control females showed a greater use of the serial strategy compared to the spatial one, potentially related to striatal function (5).

The CB1’s role in spatial information processing has been studied using a pharmaceutical approach. Interestingly, both the agonism and antagonism of CB1 impair spatial memory (*48*, *49*). By contrast, this kind of memory was improved with low doses of THC in aged mice (*50*). However, the ECS modulation of spatial memory is more complex. DAGL-KO mice with lower 2-AG concentrations had a major preference for non-spatial strategies solving MWM (*51*), while other researchers did not find any spatial memory impairment or alteration in navigation after increasing pharmacologically 2-AG or AEA levels before performing the BM test (*52*). Another possible underlying mechanism is the participation of non-classical cannabinoid receptors. For instance, the use of serial strategy was greater after applying a GPR55 agonist directly into male rats’ hippocampus, while applying an antagonist increased the use of random strategy (*53*).

Concerning sex dimorphism, it is worth highlighting that CB1 density in the brain differs significantly between sexes. Numerous studies have reported that males exhibit higher CB1 receptor density across most brain regions, while females demonstrate an enhanced receptor response following agonist stimulation (*54*, *55*). This sexual dysmorphism has also been described in pathological conditions. Early-life exposure to ketamine impairs spatial memory retrieval in adult male but not female mice, probably due to a decrease of AKTmTOR pathway (*56*). Chronic alcohol treatment in adult male mice shifted their learning from hippocampal-dependent pathways to striatum-dependent ones, which correlated to a decreased use of spatial memory (*57*). In this line, a model of alcohol binge-drinking in young mice reduced CB1 density in excitatory terminals and astrocytes in the hippocampal formation of male animals, which suggests that this shift from one specific pathway to the other could be, relatively, due to a reduced CB1 density.

In conclusion, this study highlights the crucial role of the CB1 receptor in recognition memory and the use of navigation strategies. Its involvement is both cell-type- and sex-dependent. Our data suggest that, in basal conditions, neuronal CB1 receptors in the hippocampus are necessary for expressing the spatial strategy in males but not in females. This is partly explained by the fact that females preferentially use the serial strategy over the spatial one, which is more associated with hippocampal activity. In contrast, astroglial CB1 signaling appears crucial for learning and memory processes in both sexes. Future research will explore the contributions of other cell types, such as D1+ cells, as well as the subcellular localization of CB1 and its potential sex-specific differences.

## Acknowledgements

The authors would like to thank the Animal Facility of the UPV/EHU for their support; Naroe Delgado-Martín for his assistance with the nesting analysis; and Professor Manuel Guzman for providing the *CB_1_*-flox mice.

## Funding

This work was funded by the Spanish Ministry of Science and Innovation (PGC2018-093990-A-I00 and PID2021-125763NB-I00, funded by MCIN/AEI/10.13039/501100011033 to E.S.-G.); Instituto de Salud Carlos III (PI21/00629, to S.M.; RD21/0009/0006 and RD24/0003/0027, to P.G.) and cofounded by the European Union, the Basque Government (PIBA_2023_1_0046; 2023111031; IT1473-22, to S.M.; CannaMetHD, to S.M. and G.M.; IT1620-22 to PG), ARSEP Foundation (ARSEP-1310 to S.M. and G.M.), INSERM (to G.M.), the European Research Council (MiCaBra, ERC-2017-AdG-786467, to G.M.), Fondation pour la Recherche Medicale (FRM, DRM20101220445 to G.M.), Region Aquitaine (CanBrain, AAP2022A-2021-16763610 and −17219710 to G.M.); French State/Agence Nationale de la Recherche (HippObese, ANR-23-ce14-0004-03; ERA-Net Neuron CanShank, ANR-21-NEU2-0001-04; CaMeLS, ANR-23-CE16-0022-01, to G.M.) and La Caixa Research Health 2023 (PsychoCannabis, HR23-00793, to G.M.).

M.C. was supported by the IKUR strategy grant. L.S-B. was supported by EUSKAMPUS/UPV-EHU grant.

## Notes

### Competing Interest Statement

The authors have declared no competing interest.

### Summary of Updates

- Authors' affiliations of Andres M. Baraibar - Funding from the Basque Government and Instituto Carlos III to Pedro Grandes

