## Supplementary material for "NEUROGLIAL CB1 RECEPTORS CONTROL NAVIGATION STRATEGIES": Supp.Fig 1

**Figure S1. AAVs expression in the hippocampus.** Expression of the viral vectors used in the hippocampus 7-8 weeks after injections. (A), (C) and (E) photographs show the expression of the control AAVs and (B), (D), and (F) of the CRE AAVs thanks to the mCherry tag. (A) and (B) shows the Syn-AAVs and (C) and (D) the CAMKII-AAVs (red), colocalizing with NeuN staining (green). (E) and (F) exhibits GFAP-AAVs (red) colocalizing with GFAP staining (green). Scale bars 200µm

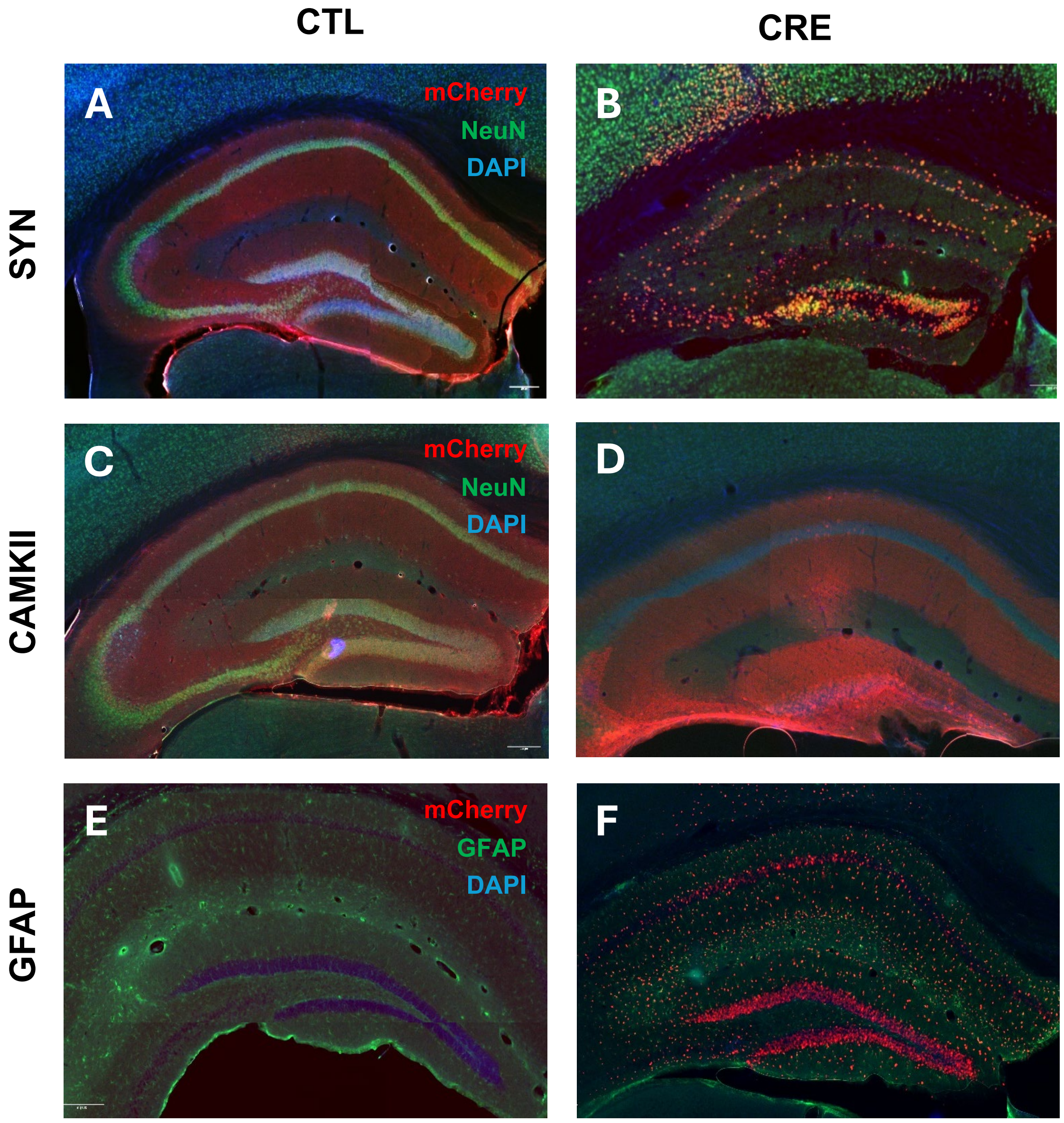
