## Supplementary material for "NEUROGLIAL CB1 RECEPTORS CONTROL NAVIGATION STRATEGIES": Supp.FIg 2

**Figure S2. CB1 deletion in the hippocampus alters the percentage of serial and random strategies.** No main differences were found in the percentage of serial **(A, left panel)** and random **(B, left)** strategies in females compared to control, except for the CB1-KO group **(A, left)**. By contrast, males with neuronal, glutamatergic, and astrocytic CB1 deletion, but not the full CB1 KO, increase the use of the random strategy **(B, right panel)**. Data are presented as mean ± s.e.m. # p<0.05 by Two-Way ANOVA test. * p<0.05, ** p<0.01 vs CTL; # p<0.05 vs CBN;  p<0.05 vs CTL;  p<0.05 vs CBN. Random strategy: Females: F (4, 38) = 1,987; ns p = 0.1161// Males: F (4, 48) = 3,589; # p = 0.0122. See **Methods** for detailed statistics.

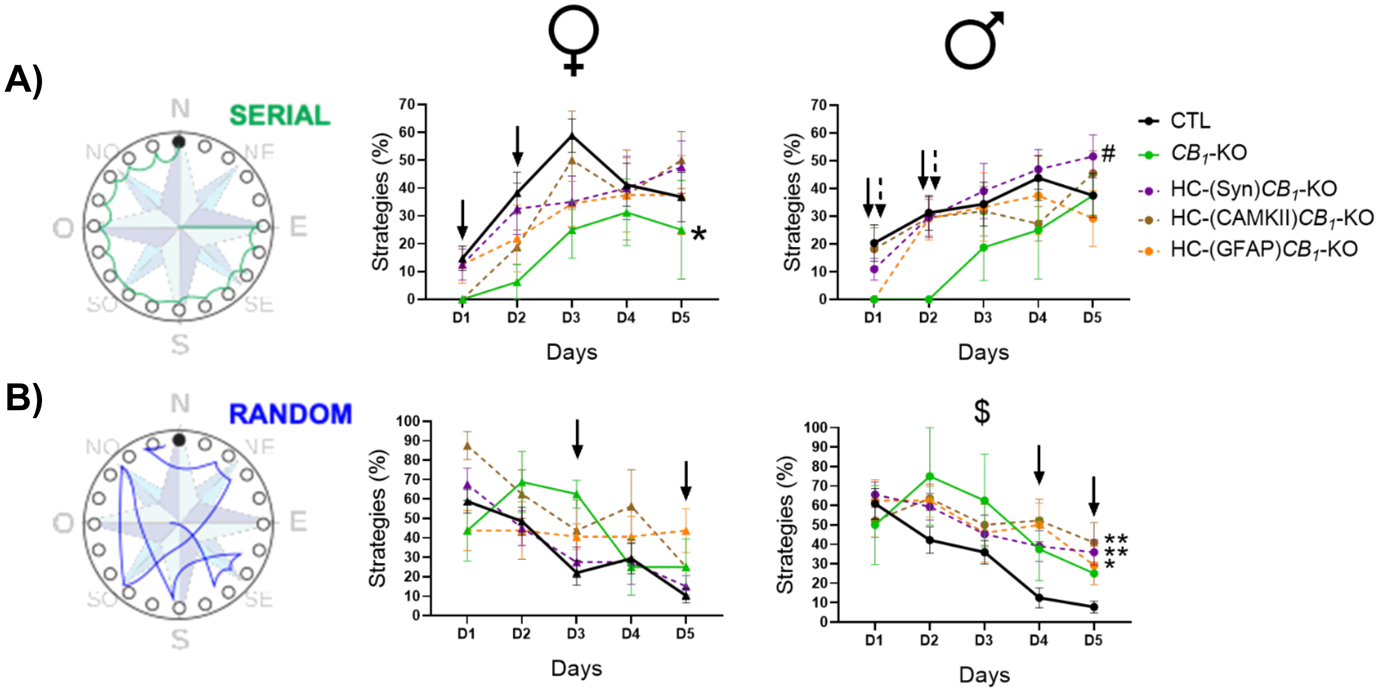
